# Implementation of the Perfect Plasticity Approximation with biogeochemical compartments in R

**DOI:** 10.1101/485128

**Authors:** Adam Erickson, Nikolay Strigul

## Abstract

Modeling forest ecosystems is a landmark challenge in science, due to the complexity of the processes involved and their importance in predicting future planetary conditions. While there are a number of open-source forest biogeochemistry models, few papers exist detailing the software development approach used to develop these models. This has left many forest biogeochemistry models large, opaque, and/or difficult to use, typically implemented in compiled languages for speed. Here, we present a forest biogeochemistry model from the SORTIE-PPA class of models, PPA-SiBGC. Our model is based on the Perfect Plasticity Approximation with simple biogeochemistry compartments and uses empirical vegetation dynamics rather than detailed prognostic processes to drive the estimation of carbon and nitrogen fluxes. This allows our model to be used with traditional forest inventory data, making it widely applicable and simple to parameterize. We detail the conceptual design of the model as well as the software implementation in the R language for statistical computing. Our aim is to provide a useful tool for the biogeochemistry modeling community that demonstrates the importance of vegetation dynamics in biogeochemical models.

## 1. Motivation and Significance

The practice of modeling ecological systems began around 1920 with a calculus model of chemical dynamics applied to the trophic interaction of herbivory [1]. The logistic growth model used was inspired by Malthusian carrying capacity [2]. The concept of *Ökologie* (ecology) had been formalized by Haeckel in 1866, with *ecosystems* soon to be coined by Tansley in 1935. Lotka-Volterra equations, a type of periodic Kolmogorov system, were next applied to fish populations [3]. Four decades later, empirical differential equations were developed to model the growth and yield of forest stands [4, 5, 6]. These models extended the principle of growth tables, used in Germany since the 18^th^ century and in China since the 17^th^ century [7].

The simplicity of early forest ecosystem models reflected the computational limits of the era – models were tractable by necessity, solved by mechanical calculators or hand. Digital computers brought a landmark innovation in the ability to explicitly simulate processes of forest succession at an individual-tree level [8]. For the first time, direct analysis of forest dynamics theory [9] was possible. These forest ‘gap’ models exhausted computational resources of the era beyond the stand or landscape scale – a limitation that continues to date. Concurrently, the first one-dimensional physiological or biogeochemical process models and forest fire models were developed. Many components of modern terrestrial biosphere models were built separately and later assembled into comprehensive global modeling systems.

Until recently, the number of transistors in integrated circuits doubled every two years in accordance with Moore’s Law. Despite the growth in compute, gap models remain computationally impractical for regional- or global-scale modeling. While many modeled processes are inherently serial, others are poised to greatly benefit from mass parallelization (e.g., using general-purpose graphics processing units, or GPGPUs). There is surprisingly little research in this area currently, as most modeling groups prefer to add new processes rather than optimize existing ones. Yet, there is a critical need to produce “a relatively simple mechanistic ecosystem model that is equitable in detail and that will run at large scales” [10]. Such a model is required to improve representation of vegetation dynamics in earth system models in order to produce more robust predictions of the global carbon cycle [11].

A lack of detailed field observations made (and still make) models of forest dynamics difficult to parameterize and validate, due to the long timescales and large number of parameters and processes involved [12]. While long-term ecological research (LTER) began formally with NSF funding of six sites in 1980, less than four decades have since passed. There are few research forests with a century of data or more. Even for sites with a long history of data (e.g., Harvard Forest), the sparsity data makes the validation of complex models non-trivial. Meanwhile, forest measurement techniques have radically advanced since the 1980s, first in the 1990s with eddy covariance flux towers and second in the 2000s with geometric point cloud models generated by laser scanning or photogrammetric computer vision (e.g., structure-from-motion). These new data sources provide detail on forest energy and biogeochemistry fluxes, canopy dynamics, species distributions, demography, and other metrics of vital importance to developing and validating new forest biogeochemistry models.

Over the past century, models of forest ecosystems grew in complexity from differential equations to detailed models of physiological and spatial processes. This progression entailed seven landmark stages of model development: (*i*) growth-and-yield tables or equations [7]; (*ii*) physical soil-plant-atmosphere continuum models [13]; (*iii*) forest fire models [14]; (*iv*) forest ‘gap’ models [8, 12]; (*v*) ‘big-leaf’ physiological process models including early land surface models [15, 16]; (*vi*) hybrid and landscape models [17, 18]; (*vii*) ‘cohort-leaf’ hybrid models including ED/ED2/FATES [19, 20], LM3-PPA [21], and the simple PPA-SiBGC compartment model presented herein. Model stages *i–v* entailed increases in complexity with each new process, resulting in the desire for new approximation schemes in stage *vii* models. While stage *vi* models expanded to include modeling spatial processes at the landscape scale, stage *vii* models blend physiological and demographic processes through robust gap model reductions. Thus, current state-of-the-art (stage *vii*) models follow the modeling approach advocated in the seminal works of [22] and [23]: *prefer realism and generality to precision*.

Models of forest ecosystems entail a number of scale- and application-specific assumptions. Historically, this has required the selection of different models for different applications or research questions [24, 25]. Terrestrial biosphere models, for example, were separated into diagnostic and prognostic models [26]. While model structure diverged over the previous four decades into specialized applications, it has converged during the past two with the development of hybrid models. The recent development of ‘cohort-leaf’ models has made this convergence complete, integrating aspects of each class of model, from individual-based gap models to global-scale terrestrial biosphere models. The distinction between diagnostic and prognostic models has similarly faded.

It is beneficial to comprehend that the mathematical approximations developed in stage VII models were made possible by detailed individual-based gap models. In effect, gap models were applied as generative models to produce data for difficult-to-measure dynamics needed in developing approximations. This is conceptually similar to the *sim2real* paradigm currently at the forefront of artificial intelligence and robotics research [27]. Using gap models as data generators was necessary due to a lack of detailed long-term observational data. In other words, the approximations are model emulators, as demonstrated in the seminal publication describing the PPA model [28, 29]. Unlike most statistical emulators (e.g., machine learning models), the PPA model is analytically tractable, thereby surpassing the requirement for an efficient model approximation providing macroscopic equations of forest dynamics [10].

In recent work [30], we demonstrated that the PPA model extended with simple biogeochemistry compartments (PPA-SiBGC) is adequate to produce model realism and precision surpassing LANDIS-II and its latest NECN biogeochemistry model, which is an adaptation of the CENTURY model. This work is important because it demonstrates that improving the representation of vegetation dynamics in forest biogeochemistry models may yield model accuracy surpassing far more complex models lacking explicit canopy dynamics. Moreover, our presented model is computationally efficient, with speeds an order of magnitude faster than LANDIS-II despite being implemented in an interpreted rather than compiled language (R rather than C#).

## 2. Conceptual Framework

In the following sections we describe the conceptual framework behind the PPA-SiBGC model.

### 2.1. Perfect Plasticity Approximation

The Perfect Plasticity Approximation (PPA) model [28] was developed based on the SORTIE individual-based model of forest ecosystems, or gap model [31]. The PPA model reduces the dimensionality of the classical SORTIE gap model by approximating the 3-D geometric interactions of individual tree crowns at the cohort level. The PPA model was based on the observation that the inclusion of phototropism (i.e., stem-leaning) and crown plasticity (i.e., space-filling) in the crown-plastic SORTIE model, CP-SORTIE, reduced the variation in canopy join height to a negligible level [28]. Thus, assuming perfect plasticity would yield zero variation in canopy join height, allowing the canopy to simply and effectively be segmented into separate canopy layers. This property is extremely important for application in modern terrestrial biosphere models, which widely adopt one-dimensional big-leaf representation of processes. The PPA model is succinctly described by Equation 1:

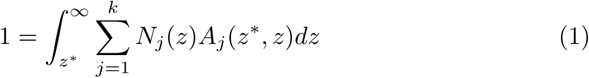

where *k* is the number of species, *j* is the species index, *N*_*j*_(*z*) is the density of species *j* at height *z, A*_*j*_(*a*^*^, *z*) is the projected crown area of species *j* at height *z*, and *dz* is the derivative of tree height. Thus, we compute the height where the integral of tree crown area is equal to the stand ground area. This yields the theoretical *z*^*^ height that marks the transition from above to below one canopy layer [28]. There may be one or many *z*^*^ heights. The number of theoretical *z*^*^ heights in a stand is a function of the stand’s leaf-area index, or LAI, where *n*_*z*\*_ = *b*LAI*c*. Each additional closed canopy layer, including shrubs and grasses, follows the form *z*^**^, *z*^***^, *et cetera*.

This partitioning of canopy layers allows for the use of separate coefficients or models of growth, mortality, and fecundity to be applied across the strata. The first moment of these canopy layer dynamics accurately approximates the dynamics of individual-based models [28]. We extend the SORTIE-PPA model by adding a simple compartment-based representation of biogeochemistry using allometric and stoichiometric relations, along with simple prognostic (i.e., climate-driven) model of soil respiration [32, 33] and a constant representation of organic carbon by soil type [34].

### 2.2. Allometry and Stoichiometry

The tree allometric model and parameters were adapted from previously published research [35, 36]. Tree height is modeled as a non-linear function of stem diameter as follows:

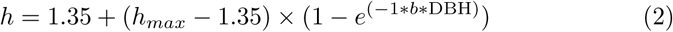

where *h* is the tree height (m), *h*_*max*_ is the maximum potential tree height, DBH is the depth-at-breast-height (cm), *e* is Euler’s constant, and *b* is an exponential decay coefficient. Tree crown radius and depth are also modeled as a function of DBH, but are instead intercept-free linear models.

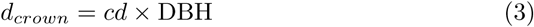

where *dp*_*crown*_ is the crown depth (m) and *cd* is the crown depth regression coefficient. The equation form is similar for crown radius:

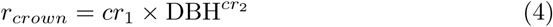

where *r*_*crown*_ is the crown radius (m) and, *cr*_1_ and *cr*_2_ are the crown radius regression coefficients. Basal area is also calculated as a function of DBH, using the forester’s constant for DBH in centimeters:

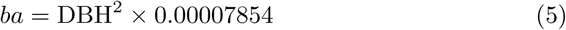

National species-specific biomass equations [37] were used to model tree biomass as a function of DBH:

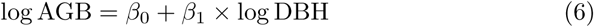

where AGB is the tree aboveground biomass (kg) and,; *β*_0_ and; *β*_1_ are regression coefficients. Empirical coefficients are used for the aboveground biomass fractions contained in stem, branch, and leaf compartments, as well as soil. Root biomass is partitioned into coarse and fine root components based on existing equations for the United States [37], following the general form:

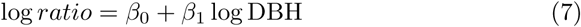

where *ratio* is the biomass fraction for the root component and, *β*_0_ and *β*_1_ are regression coefficients. The coefficients used for coarse roots were −1.4485 for *β*_0_ and −0.03476 for *β*_1_. For fine roots, the coefficients were −1.8629 and −0.77534, respectively. The biomass of each root compartment is then calculated by multiplying tree AGB by the corresponding ratio.

Tree belowground biomass is calculated as the sum of root and soil biomass, while total biomass is calculated as the sum of below- and above-ground compartments. Separate empirical biomass carbon fraction and C:N stoichiometric coefficients were used for each compartment. Thus, C and N content are fixed fractions of biomass values, based on empirical point estimates or samples from distributions.

### 2.3. Soil Respiration

In the PPA-SiBGC model, we use the simple prognostic soil respiration model of Raich *et al.* (2002):

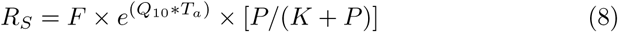

where *e* is Euler’s constant, *Q*_10_ is the respiration temperature sensitivity coefficient per 10 °C increase, *b* is a temperature sensitivity constant, *T*_*a*_ is mean monthly air temperature (°C), *P* is mean monthly precipitation (cm), *F* is the soil respiration rate at 0 °C, *K* is the half-saturation constant for the hyperbolic relationship between soil respiration and rainfall, and *R*_*S*_ is soil respiration (g C m^2^ day^−1^). The version of the model used in PPA-SiBGC includes updates parameters based on infrared gas analyzer measurements of soil CO_2_ flux [33].

### 2.4. Soil Organic Carbon

For modeling soil organic carbon (SOC) in PPA-SiBGC, we use the simple approach of Domke *et al.* (2017), which is based on the STATSGO US national soil database. Thus, the model is currently limited to forest soil types present in the US. The model is defined as follows:

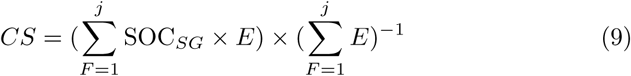

where *CS* is the county-specific areal SOC by forest type (Mg/ha), *SOC*_*SG*_ is the areal SOC for the STATSGO map unit (Mg/ha), *E* is a vector of weights for the areal coverage of each USFS Forest Inventory and Analysis (FIA) plot, and *F* is the number of FIA plot records within forest types. We used the best fit model of [34] to model the vertical SOC profile:

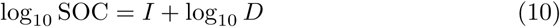

where log_10_ SOC is the volumetric soil organic carbon density (Mg C ha^−1^cm^−1^), *I* is the intercept, and *D* is the profile midpoint depth (cm). We integrated over a range of profile depths to produce total SOC values:

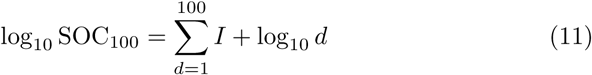

where *d* is the profile midpoint depth (cm). And thus:

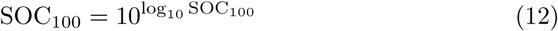

Both the original SORTIE-PPA model and the PPA-SiBGC model presented herein have undergone extensive validation with field data. The SORTIE-PPA model validation is described in the original paper [28], while the PPA-SiBGC model was recently validated at two research forests for a range of metrics [30]. Despite the simplicity of the representation of biogeochemical dynamics in the PPA-SiBGC model, it outperformed LANDIS-II with NECN (i.e., CENTURY) biogeochemistry across a range of metrics and sites in the model intercomparison exercise [30].

## 3. Software Description

The PPA-SiBGC model is implemented in a standalone R script that is designed to be run from a command-line interface, or CLI. Since the model is implemented in R with no external dependencies, it can be used on any platform that R supports (e.g., Windows, Linux, MacOS). The model implementation begins by loading all input data into memory and parsing the configuration CSV file. If cohort mode is enabled, the *Generate Cohorts* function aggregates the individual trees into cohorts based on a predefined DBH interval (e.g., every 2 cm), recording the number of trees per cohort.

Next, the allometric and stoichiometric C:N equations are applied to the individual trees or cohorts in order to calculate the initial C and N pools. The simulation is then run for each year in the defined temporal range. The overall model process is shown in Algorithm 1.

### Algorithm 1 PPA-SiBGC model procedure

~~~
1: **procedure** MODEL
2:   Load input files
3:   Generate cohorts
4:   Initialize tree allometry and biogeochemistry
5:   Calculate soil organic matter profile
6:   **for** year, …, *N*_*years*_ **do**
7:      Calculate soil respiration
8:      Calculate *z*^*^ height (PPA algorithm)
9:      **for** species, …, *N*_*species*_ **do**
10:      **for** type, …, *N*_*types*_ **do**
11:          Apply mortality
12:          Apply growth
13:          Calculate allometry
14:      **end for**
15:      Calculate biomass
16:      Calculate C and N
17:   **end for**
18:   Append annual outputs to CSV
19:   Calculate year execution time
20:  **end for**
21:  Calculate total execution time and export to CSV
22: **end procedure**
~~~

Vectorization is used where possible to accelerate the model operations. This optimization comes at no cost to the programmer in interpreted languages such as R, as it is built into the language. Distributions such as Microsoft R Open (MRO) ship with the Intel MKL optimized algebra library. An implementation of the PPA algorithm used to find the theoretical *z*^*^ height is shown in Algorithm 2:

### Algorithm 2 simplified PPA algorithm

~~~
**Input:** *T*_1_ *… T*_*N*_ (tree list from forest inventory), *A*_*field*_ (field area)
**Output:** *T* (height-sorted tree list), *z*_*star*_ (calculated *z*^*^ height)

1: **procedure** PPA(*T*, *A*_*Field*_)
2:    *z*_*Star*_ ← *NULL*
3:    *T* ← Sort(*T*_*Height*_, *Descending*)       ⊲ sort trees by descending height
4:   **for** i = 2, …, *T*_*N*_ **do**
5:        *CrownArea*_*T*_ [*i*] ← *CrownArea*_*T*_ [*i*] + *CrownArea*_*T*_ [*i* –1]
6:        **if** *CrownArea*_*T*_ [*i*] > *A*_*Field*_ **then**
7:            *z*_*Star*_ ← *Height*_*T*_ [*i*]
8:            **return** *T*, *z*_*Star*_             ⊲ return from function
9:        **else**
10:           *continue*
11:       **end if**
12:  **end for**
13:  **return** *T*, *z*_*Star*_                 ⊲ return from function
14: **end procedure**
~~~

The software implementation provided is an approximation of the PPA algorithm that simplifies its calculation:

~~~
Trees <- Trees [ **order** (Trees**$** height, decreasing=TRUE), ]
Trees**$**canopy_area <- **cumsum** (Trees**$**crown_a **\*** Trees**$**n_trees)
**index** <- **which**. **min** (**abs** (Trees**$**canopy_area - field_area))
zstar <- Trees [ **index**,]
~~~

Our implementation of the soil respiration model [33] is straightforward, as shown below:

~~~
respiration_soil <- **function** (Ta, P) {
  **if** (Ta < –13.3) {
   Rs = 0
} **else** {
   **if** (Ta > 33.5) { Ta = 33.5 }
   e = **exp** (1)
   Q10 = 0. 05452 (C −1)
   F = 1. 250
   K = 4. 259
   Rs = F **\*** ê (Q10 **\*** Ta) **\*** (P **/** (K + P))
  }
   **return** (Rs)
}
~~~

Meanwhile, our implementation of the soil organic carbon (SOC) profile model [34] is based on a lookup table containing soil classes and corresponding regression model intercepts and slopes. The parameters are extracted and the linear model is applied to calculate SOC along the defined profile interval. By default, the profile interval is set to [1 .. 100].

~~~
soc_depth <- **function** (**order**, depth_**cm**=100) {
  soc_**table** = **data**. **frame** (
    **order** = **c** (” All ”, ” Alfisols ”, ” Andisols ”,
    ”Aridisols ”, ” Entisols ”, ” Histosols”, ”Inceptisols”,
    ”Mollisols”,”Soosols”,”Ultisols”,”Vertisols”),
    intercept = **c**(1.1795,1.1122, 1.3837,0.2065,0.9300,
    1.6227, 1.1631, 1.0163,1.4262, 1.1576,0.5145),
    slope = **c**(-0.8228, -0.8330, -0.8425, -0.1300,
    -0.7207, -1.0109, -0.7331, -0.6214, -0.9801, -0.8867,
    -0.2427)
  )
  rowval = soc_**table** [ soc_**table** \**$order**==**order**,]
  coeffs = **as**. **numeric** (rowval [, **c** (” intercept ”, ” slope ”) ])
  soc = **sapply** (**seq** (1, depth_**cm**, 1), **function** (x) *{*
  10^ (coeffs [ 1 ] + coeffs [ 2 ] **\* log10** (x))
  })
  **return** (**sum**(soc))
}
~~~

## 4. Illustrative Example

For model parameterization, validation, and comparison with the LANDIS-II model at Harvard Forest EMS flux tower in Massachusetts, USA and Jones Ecological Research Center RD flux tower in Georgia, USA, readers may refer to our recent model intercomparison paper [30]. An example of running the PPA-SiBGC program (ppa v50.r) from the CLI is shown below:

~~~
Rscript --vanilla ppa_v50. r --wd /home/ model --verbose
~~~

Here, the *Rscript* executable is used with the *–vanilla* option to run the program within a new R session. Additional options include the *–wd* flag to specify a working or target directory containing parameter files in a pre-defined directory structure and the *–verbose* flag to run the model in verbose mode for monitoring progress or debugging. The predefined directory structure expected for model input files containing parameters and drivers (i.e., climate data) is as follows:

**Figure 1:**
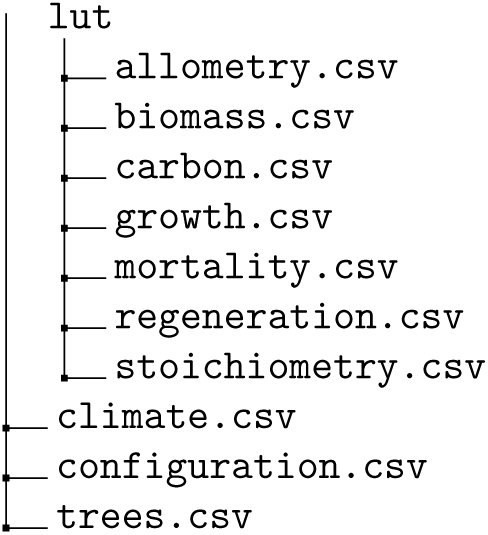
Directory structure of PPA-SiBGC inputs.

When the simulation run is completed, the following CSV files are produced in an *outputs* directory within the target directory:

**Figure 2:**
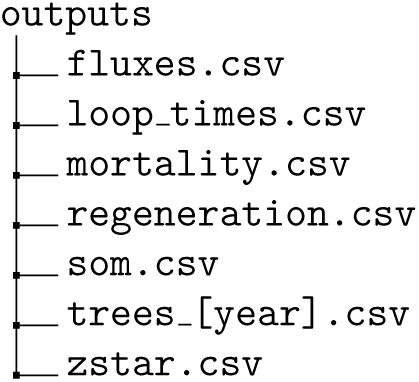
Directory structure of PPA-SiBGC outputs.

These outputs contain the ecosystem biogeochemistry fluxes, time to complete each model iteration, cohort mortality with allometry and biogeochemistry, species regeneration, soil organic matter (SOM) pools, cohort list by simulation year with allometry and biogeochemistry, and the theoretical *z*^*^ height by year.

## 5. Anticipated Impact

We anticipate that the PPA-SiBGC model will showcase the importance of realistically approximating vegetation dynamics in order to develop forest biogeochemistry models with improved generality and precision. This is because the inclusion of even simple allometric and stoichiometric biogeochemistry relations with the PPA model showed accurate estimation of ecosystem fluxes [30]. The robustness of allometric scaling theory, rooted in the self-similarity of tree species and shaped by physical constraints, is well supported in theoretical research using highly detailed models [38]. Thus, species-specific allometric models remain a useful modeling abstraction in global-scale modeling.

We believe that our work also demonstrates the the classical modeling trade-off of Levins (1966) between generality, precision, and realism is unduly imposed; in our work, model generality and precision were possible only through enhanced model realism. Thus, no trade-off was apparent in our case. We therefore reiterate his suggestion that modelers focus on improving realism over generality and precision, which may result in improvements of all three criteria.

From a practical standpoint, we hope that our work will help to advance the field by making forest biogeochemistry models more approachable, cross-platform, and easier to use. All input and output files use a CSV file structure in order to facilitate simplified key-value parsing of all data in user-developed programs. This is in contrast to models such as LANDIS-II that use unstructured TXT files that are laborious and inefficient to parse. In future work, we will soon release model wrapper libraries in R and Python for simplifying the operation, parameterization, optimization, and validation of forest ecosystem models.

## 6. Conclusions

In conclusion, we provide a new forest biogeochemistry model, PPA-SiBGC, based on the Perfect Plasticity Approximation (PPA) algorithm in the R language of statistical computing. The program is cross-platform and is designed to be simple to deploy and apply. The program is designed to run from a command-line interface, while all model inputs and outputs are in CSV format to facilitate simplified data pre- and post-processing. The structure of the program is simple and effective, using vectorization where possible to speed operations without the programmer and computational overhead of multi-thread or multi-core parallelization. Our model uses only base R libraries, facilitating ease of deployment across a variety of systems. We demonstrate that effective forest biogeochemistry models need not be comprised of hundreds of thousands of lines of code in difficult-to-use compiled languages. In future work, we will release forest biogeochemistry model wrapper libraries in R and Python to ease the operation and extend the use of this and other forest biogeochemistry models. All model code is made available at our GitHub repository under an Apache 2.0 license:

https://github.com/adam-erickson/ecosystem-model-comparison/blob/master/models/ppa_bgc/hf_ems/ppa_v50.r

## Acknowledgements

Funding: This work was supported by the U.S. Army Corps of Engineers [contract number W912HQ-18-C-0007].

